# Tasmanian devil facial tumor-derived extracellular vesicles reveal mesenchymal transition markers and adhesion molecules related to metastasis

**DOI:** 10.1101/2020.10.18.344721

**Authors:** Camila Espejo, Richard Wilson, Gregory M. Woods, Eduard Willms, Andrew F. Hill, A. Bruce Lyons

**Affiliations:** Tasmanian School of Medicine, College of Health and Medicine, University of Tasmania, Hobart, TAS 7000, Australia; Central Science Laboratory, University of Tasmania, Hobart, TAS 7005, Australia; Menzies Institute for Medical Research, College of Health and Medicine, University of Tasmania, Hobart, TAS 7000, Australia; Department of Biochemistry and Genetics, La Trobe Institute for Molecular Science, La Trobe University, Bundoora, VIC, 3083, Australia

## Abstract

Tasmanian devils are threatened with extinction by Devil Facial Tumor Disease (DFTD), which consists of two genetically independent transmissible cancers (DFT1 and DFT2). Both cancers typically cause death due to metastases. However, the mechanisms underpinning DFTD metastasis are not well understood. The nano-sized, membrane-enclosed extracellular vesicles (EVs) released by cancer cells have been implicated in metastasis, thus EVs may yield insights into DFTD metastasis. Here, we characterized EVs derived from cultured DFT1, DFT2, and devil fibroblast cells. The proteome of EVs was determined using data independent acquisition mass spectrometry and an in-house spectral library of >1,500 proteins. Relative to EVs from fibroblast cells, EVs from both DFT1 and DFT2 cell lines expressed higher levels of proteins associated with cell adhesion and focal adhesion functions. Furthermore, hallmark proteins of epithelial mesenchymal transition, which are associated with increased metastatic features in some cancers, were enriched in DFT2 EVs relative to DFT1 EVs, suggesting differential aggressiveness between the cancers and a target for novel differential diagnosis biomarkers. This first EV-based investigation of DFTD increases our understanding of the cancers’ EVs and their possible involvement in the metastatic process. As EVs are found in body fluids, these results offer potential for non-invasive biomarkers for DFTD.

## Introduction

The Tasmanian devil (Sarcophilus *harrisii*) is the world’s largest extant carnivorous marsupial, currently threatened with extinction due to two transmissible cancers causing Devil Facial Tumor Disease (DFTD). The first described DFTD, (referred to as DFT1), has dramatically impacted devil numbers, which have declined by 77% since DFTD was discovered in 1996 (1). DFT1 is caused by a clonal cell of Schwann cell origin (2), transmitted among devils when transplanted through biting during social interactions (3). Fatal in almost 100% of cases, DFT1 kills its host within 6 to 12 months after the presentation of tumor masses on facial, oral and neck regions (4, 5).

The second genetically independent transmissible cancer (DFT2) was first reported in 2014 and is also of Schwann cell origin (6). Unlike DFT1, DFT2 appears to have originated in a male individual (6) and while DFT1 has widely spread across Tasmania since its discovery (7), DFT2 is currently confined to the Channel region of Southeast Tasmania. Death of the host has been proposed to be a consequence of three main mechanisms: metastasis, starvation and secondary complications (8). With DFT1, metastasis occurs in approximately 65% of cases, primarily affecting lungs and spleen (5). However, the timing and mechanisms of DFT1 metastasis require further investigation to provide a better understanding of the pathogenesis and progression of the cancer.

DFT1 and DFT2 are grossly indistinguishable but can be differentially diagnosed using histopathology, immunohistochemistry, cytogenetics, and PCR techniques (6, 9, 10). These laboratory techniques require time and expertise, and the collection of a tumor biopsy from the infected animal. Development of a rapid non-invasive differential diagnostic test for DFT1 and DFT2 would significantly improve the potential scope and scale of disease surveillance.

Most cells, including cancer cells in humans and animal models, secrete extracellular vesicles (EVs) into their extracellular environment. EVs are lipid bilayer structures which contain bioactive cargo, such as proteins, genetic material, and lipids (11). Some EV cargo represent the proteome of the parent cell, but may also be enriched in a subset of common EV marker proteins related to their biogenesis (12). EVs have been proposed to be major players in the process of intercellular communication, through the transfer of their cargo, both locally and systemically (13). Using transcellular and paracellular routes, EVs can enter a variety of body fluids, including blood (14). The EV phospholipid bilayer membrane protects cargo against degradation by serum proteases and nucleases (15). These properties have made EVs an increasingly attractive tool for diagnostic biomarker discovery in cancer research and other diseases (16).

A major hallmark of cancer is that tumor cells need to communicate with other cells to survive, invade and progress. As EVs play a crucial role in intercellular communication, they are active participants in the process of tumorigenesis and cancer progression (17). Thus, EVs originating from tumor cells, or the tumor microenvironment, can support tumor cell growth and promote successful colonization of local and distant organs (18–20). EVs facilitate the process of metastasis through a variety of mechanisms including immune modulation, microenvironment remodeling, angiogenesis, intravasation and extravasation of tumor cells, and preparation of the metastasis niche at the target organs (21, 22). As metastasis is the leading cause of death in cancer patients (23), and EVs express molecules which are associated with cancer progression, they have been investigated as a source of prognostic biomarkers for cancer (24).

Here, we provide the first analysis of the proteome of EVs from DFTD cells. We identified potential metastasis mechanisms and proposed a potential application of EVs for prognosis and differential diagnosis of DFTD tumors.

## Experimental procedures

### Cell culture

Three DFTD cells lines and healthy devil fibroblast were cultured in RPMI medium (Gibco no. 11875-093), supplemented with 10% heat-inactivated fetal bovine serum (FBS), and 1% antibiotic-antimycotic (Thermo Scientific no. 15240062). Each cell line was cultured in duplicates in a 175 cm^2^ culture flask (Corning) at 35°C in a fully humidified atmosphere of 5% CO_2_.

### Isolation of extracellular vesicles

Cell cultures were used at 60-70% confluence. Culture medium was discarded, and cells were washed twice with phosphate-buffered saline (PBS) and then incubated in culture medium, supplemented with 5% of heat-inactivated FBS, for 48 hours. This medium had been previously subjected to centrifugation at 100,000 *g* for 18 hr at 4 °C to deplete bovine EVs from FBS (25). After 48 hours, the cultured medium of each cell line was collected and centrifuged at 1,500 *g* for 10 minutes at 4 °C to remove cells and debris. The supernatant was further clarified by centrifugation at 10,000 *g* for 10 minutes at 4 °C. Supernatants were then concentrated using Amicon Ultra-15 centrifugal filters (MWCO 100 kDa) to a final volume of 2 ml. Then following the manufacturer’s instructions, the concentrated supernatant was subjected to size exclusion chromatography on qEV2 – 35 columns (IZON). Briefly, EVs were eluted in PBS containing 0.05% sodium azide in eight fractions of 1 ml each were eluted after a 14 ml void volume and pooled. The EV samples were concentrated with Amicon Ultra-15 centrifugal filters (MWCO 100 kDa) to a final volume of 500 μl and stored at −80 °C until further use.

### Electron microscopy

The EVs were imaged using a JEM – 2100 transmission electron microscope (JEOL Tokyo, Japan) equipped with a LaB6 filament operating at 200 kV. Images were recorded using a Gatan Orius SC200 2k x 2k charge-coupled device (CCD) camera at a range of magnifications. 400-mesh carbon coated copper grids (ProSciTech) which had been glow-discharged in air to render them hydrophilic using an Emitech 950X equipped with a K350 glow discharge unit (Quorum Technologies) were used. 10 μl of the isolated EV samples obtained with the SEC columns were dropped onto the prepared grids and left for at least 30 seconds. Excess fluid was drawn off with filter paper, two drops of 2% Uranyl Acetate were added for approximately 10 seconds each before being drawn off with filter paper. The grids were then dried and transferred into the TEM for viewing.

### Nano particle tracking analysis (Nanosight)

EV size distribution and concentration were determined using a Nanosight NS300 nanoparticle analyzer (Malvern Panalytical, Malvern, UK) equipped with a 405 nm laser and Nanosight NTA 3.2 software. Samples were measured in PBS, and camera level was set at 14 for all recordings. Camera focus was adjusted to make the particles appear as sharp individual dots, and three 30-second videos were recorded for each sample. For post-acquisition analysis, all functions were set to automatic except detection threshold, which was set at 5. EV size data were normalized to the equal area under the curve for comparison between samples.

### Protein preparation

EV proteins were extracted according to the method of Abramowicz, et al. (26). EV samples were mixed with acetonitrile to a final concentration of 50% (v/v) and evaporated using a centrifugal vacuum concentrator. Protein samples were resuspended in 100 μl of denaturation buffer (7M urea and 2 M thiourea in 40 mM Tris, pH 8.0).

Duplicate samples of the cell lines were washed twice in PBS after collection of cultured supernatants for EVs isolation. A cell count of 1×10^7^ cells per ml was determined using a hemocytometer. Pelleted cells were carefully lysed in 700 μl of denaturation buffer supplemented with 1% (w/v) of Halt protease inhibitor cocktail (100X, Thermo Scientific) and then incubated on a tube rotator for 2 hrs at 4 °C. The lysate was centrifuged at 13,000 *g* for 10 minutes and the supernatant finally cleaned up by precipitation using 9x volumes of 100% ethanol overnight at −20 °C. Precipitated proteins were pelleted by centrifugation at 13,000 *g* for 10 minutes. Protein pellets were briefly air-dried and then reconstituted in 100 μl of denaturation buffer.

Protein concentration from lysates and EV samples was determined using EZQ protein quantification Kit (Thermo Scientific). For mass spectrometry analysis, aliquots of 30 μg of protein were sequentially reduced using 10 mM DTT overnight at 4 °C, alkylated using 50 mM iodoacetamide for 2 hrs at ambient temperature and then digested with 1.2 μg proteomics-grade trypsin/LysC (Promega) according to the SP3 protocol (27). Digests were acidified by the addition of TFA to 0.1% and peptides collected by centrifugation at 21,000 x g for 20 minutes. Samples were further cleaned up by offline desalting using ZipTips (Merck) according to manufacturer’s instructions.

### SDS-PAGE and Western blotting

EVs and lysate samples resuspended in denaturation buffer were mixed with freshly β-mercaptoethanol (5% (v/v)) (Sigma) and heated for 10 minutes at 95 °C. Protein samples (20 μg in each lane) were separated on a Bolt™ 4 - 12%, Bis-Tris, 1.0 mm, Mini Protein Gel (Thermo Scientific Invitrogen™) in NuPAGE™ MES SDS Running Buffer, alongside a molecular weight marker (SeeBlue™ Plus2 Pre-stained Protein Standard, Thermo Scientific Invitrogen™). Blotting was performed on an Immobilin®-P PVDF membrane with 0.45 μm pore size (Merck Millipore) using a Mini Blot Module (Thermo Scientific Invitrogen™).

Membranes were blocked in 5% skimmed milk in TBS containing 0.1% Tween-20 (TBS-T) for 1 hour at room temperature, primary antibodies rabbit anti-Syntenin - ab19903 (Abcam) at 1:1000, mouse anti-Flotillin-1 - BD610821 (BD Transduction Laboratories) at 1:1000, and mouse anti-Golgi matrix protein - BD610822 at 1:1000 (BD Transduction Laboratories) were incubated overnight in 5% skimmed milk in TBS-T at 4 °C. After incubation, membranes were washed three times in TBS-T for 10 minutes at room temperature. Membranes were subsequently probed with Amersham ECL Mouse IgG, HRP-linked whole Ab (from sheep) (NA931) or Amersham ECL Rabbit IgG, HRP-linked whole Ab (from donkey) (NA934) at 1:5000 in TBS-T for 1 hour at room temperature. After incubation, membranes were washed three times in TBS-T for 10 minutes at room temperature, and subsequently visualized using Clarity Western ECL Substrate (Bio-Rad). Images were acquired on a ChemiDoc™ Touch Imaging System (Bio-Rad) and analyzed with Image Lab Software (Bio-Rad).

### Liquid chromatography and mass spectrometry analysis

#### High-pH peptide fractionation

Two experiment-specific peptide spectral libraries were generated for cell lysates and cell culture using offline high-pH fractionation. For each library, a pooled digest comprising aliquots of each sample set (180 μg) was desalted using Pierce desalting spin columns (Thermo Scientific) according to the manufacturer’s guidelines. Each sample was evaporated to dryness then resuspended in 25 μl HPLC loading buffer (2 % acetontrile containing 0.05 % TFA) and injected onto a 100 × 1 mm Hypersil GOLD (particle size 1.9 μm) HPLC column. Peptides were separated using an Ultimate 3000 RSLC system equipped with microfractionation and automated sample concatenation, operated at 30 μl/minute using a 40 minute linear gradient of 96% mobile phase A (water containing 1% triethylamine, adjusted to pH 9.6 using acetic acid) to 50% mobile phase B (80% actetonitrile containing 1% triethylamine), followed by 6 minutes washing in 90% B and re-equilibration in 96% A for eight minutes. Sixteen concatenated fractions were collected into 0.5 ml low-bind Eppendorf tubes, evaporated to dryness then reconstituted in 12 μl HPLC loading buffer.

#### Mass spectrometry – data-dependent acquisition (DDA)

Peptide fractions were analysed by nanoflow HPLC-MS/MS using an Ultimate 3000 nano RSLC system (Thermo Scientific) coupled with a Q-Exactive HF mass spectrometer fitted with a nanospray Flex ion source (Thermo Scientific) and controlled using Xcalibur software (ver 4.3). Appoximately 1 μg of each fraction was injected and separated using a 90-minute segmented gradient by preconcentration onto a 20 mm × 75 μm PepMap 100 C18 trapping column then separation on a 250 mm × 75 μm PepMap 100 C18 analytical column at a flow rate of 300 nL/min and held at 45°C. MS Tune software (version 2.9) parameters used for data acquisition were: 2.0 kV spray voltage, S-lens RF level of 60 and heated capillary set to 250°C. MS1 spectra (390 - 1500 *m/z*) were aquired at a scan resolution of 60,000 followed by MS2 scans using a Top15 DDA method, with 20-second dynamic exclusion of fragmented peptides. MS2 spectra were acquired at a resolution of 15,000 using an AGC target of 2e5, maximum IT of 28ms and normalized collision energy of 30.

#### Mass spectrometry - data-independent acquisition (DIA)

Individual peptide samples were analysed by nanoflow HPLC-MS/MS using the instrumentation and LC gradient conditions described above but using DIA mode. The sequence of sample injections was randomized by blinding the MS operator to the sample codes. MS1 spectra (390 - 1240 *m/z*) were acquired at 60,000k resolution, followed by sequential MS2 scans across 26 DIA x 25 amu windows over the range of 397.5-1027.5 *m/z*, with 1 amu overlap between sequential windows. MS2 spectra were acquired at a resolution of 30,000 using an AGC target of 1e6, maximum IT of 55 ms and normalized collision energy of 27.

#### Raw data processing

Both DDA-MS and DIA-MS raw files were processed using Spectronaut software (version 13.12, Biognosys AB). Each project-specific library was generated using the Pulsar search engine to search DDA MS2 spectra against the *Sarcophilus harrisii* UniProt reference proteome (comprising 22,388 entries, November 2016). With the exception that single-hit proteins were excluded, default (BGS factory) settings were used for both spectral library generation and DIA data extraction. For library generation, these included N-terminal acetylation and methionine oxidation as variable modifications and cysteine carbamidomethylation as a fixed modification, up to two missed cleavages allowed and peptide, protein and PSM thresholds set to 0.01. Mass tolerances were based on first pass calibration and extensive calibration for the calibration and main searches, respectively, with correction factors set to 1 at the MS1 and MS2 levels. The parameters used for DIA-MS data analysis are provided in full in the BGS setup files (supplemetal file 3 and 4). Briefly, XIC extraction deployed dynamic retention time alignment with a correction factor of 1. Protein identification deployed a 1% q-value cut-off at precursor and protein levels, automatic generation of mutated peptide decoys based on 10% of the library and dynamic decoy limitation for protein identification. MS2-level data were used for relative peptide quantitation between experimental samples, using the intensity values for the Top3 peptides (stripped sequences) and cross-run normalization based on median peptide intensity.

#### Statistical analysis

Spectronaut protein quantitation pivot reports, including protein description, gene names and UniProt accesion numbers, were uploaded into Perseus software (version 1.6.10.50) for further data processing and statistical analysis. Quantitative values were log2 transformed and proteins filtered according to the number of valid values. Proteins detected in <50% samples were excluded from EV and lysate datasets. Remaining missing values were imputed with random intensity values for low-abundance proteins based on a normal abundance distribution using default Perseus settings. Principal component analysis (PCA) was carried out using the filtered proteins for the EV and lysate samples in order to reduce the dimensionality of the dataset. Differential abundance of proteins between sample groups was determined using Student’s *t*-test with a permutation-based false discovery rate (FDR) controlled at 5% and s0 values set to 0.1 to exclude proteins with very small differences between means. Hierarchical clustering was used to group significant proteins into clusters, based on the subset of proteins that were significant according to one-way ANOVA using FDR-based truncation (5%).

#### Bioinformatics analysis

FUNRICH version 3.1.3 (28) was used to compare the EV proteins derived from the Tasmanian devil cell lines with the human Vesiclepedia, and with the top 100 exosomal proteins reported in the Exocarta database (29). We compared the proteome of cell lysates and EVs, and used DAVID software for functional enrichment analysis of the proteins present only in EVs (30, 31). DAVID software was also used for bioinformatic analysis of proteins that were upregulated in both tumor cell EVs and lysates vs. fibroblast EVs and lysates to identify whether DFTD EVs maintain key features of their cell of origin.

We combined the proteome of EVs derived from DFT1 and DFT2 cell lines for functional enrichment analysis of proteins in EVs from both cancers relative to devil fibroblast derived EVs. For this analysis we used two different bioinformatic approaches. Over Representation Analysis (ORA) is used to determine whether particular biological functions or processes are over-represented in a subset of genes or proteins when compared to a background proteome (32). We used ORA to analyze the subset of proteins demonstrating a pattern of upregulation in EVs derived from DFTD cells vs. EVs derived from devil fibroblasts determined by hierarchical cluster analysis. UniProt accessions for these proteins were uploaded to DAVID (30, 31) to identify clusters of GO terms, protein families and pathways. Functional terms with p-values < 0.05 after Benjamini-Hochberg correction for multiple hypothesis testing were considered significant (30). The protein list was analyzed using the *Sarcophilus harrisii* species proteome UniProt database. To complement the ORA of the subset of proteins that were significantly enriched and upregulated in DFTD derived EVs vs EVs derived from devil fibroblasts, we performed Gene Set Enrichment Analysis (GSEA) to analyze patterns across the entire post-filtering EV proteome. GSEA tests for non-random patterns of enrichment of functional protein groups with quantitative expression data rather than just presence of proteins names within a subset, and thus can detect subtle changes in biological functions evidenced in coordinated way in a set of related genes (32). Additionally, we performed GSEA functional enrichment analysis of EV proteins derived from DFT1 vs. DFT2 cell lines. GSEA software ver. 4.0.3 (33, 34) combined with the Enrichment map application in Cytoscape software (ver. 3.8.0) was used for network-based visualization for GSEA results, according to the method of Reimand, et al. (35). Clusters of similar pathways were automatically defined and summarized using the AutoAnnote Cytoscape application (36).

### Experimental design and statistical rationale

Technical duplicates of EVs and cell lysates from tumor cells lines obtained from three devils with confirmed DFT1 (C5065, 1426 and 4906), three devils with confirmed DFT2 (JV, SNUG and RV) were subjected to mass spectrometry analyses. Technical duplicates of EVs and cell lysates from short term fibroblasts cultures derived from three healthy devils were used as controls. Proteome from cell lysates were characterized to compare with the EV proteome, and thus to determine the extent to which the cargo of EVs reflects the parent cell. Statistical analysis was performed as described above.

## Results

### Characterization of EVs from Devil Facial Tumour (DFTD) and devil fibroblast cells

Transmission electron microscopy revealed the morphology of EVs as small closed vesicles with a cup shaped structure, consistent with previous analysis (37) (Fig. 1A). Nanoparticle tracking analysis (NTA) demonstrated successful isolation of particles with a typical small/medium EV size-distribution (38). The mean ± standard deviation diameters were 185.9 ± 78.6 nm for DFT1 EVs, 217 ± 91.2 nm for DFT2 EVs and 183.2 ± 78.6 nm for devil fibroblast EVs (Fig. 1B). Additionally, NTA demonstrated that DFT2 EVs were secreted in significantly higher numbers than EVs derived from DFT1 and devil fibroblast cell lines (Fig. 1C).

**Figure 1.**
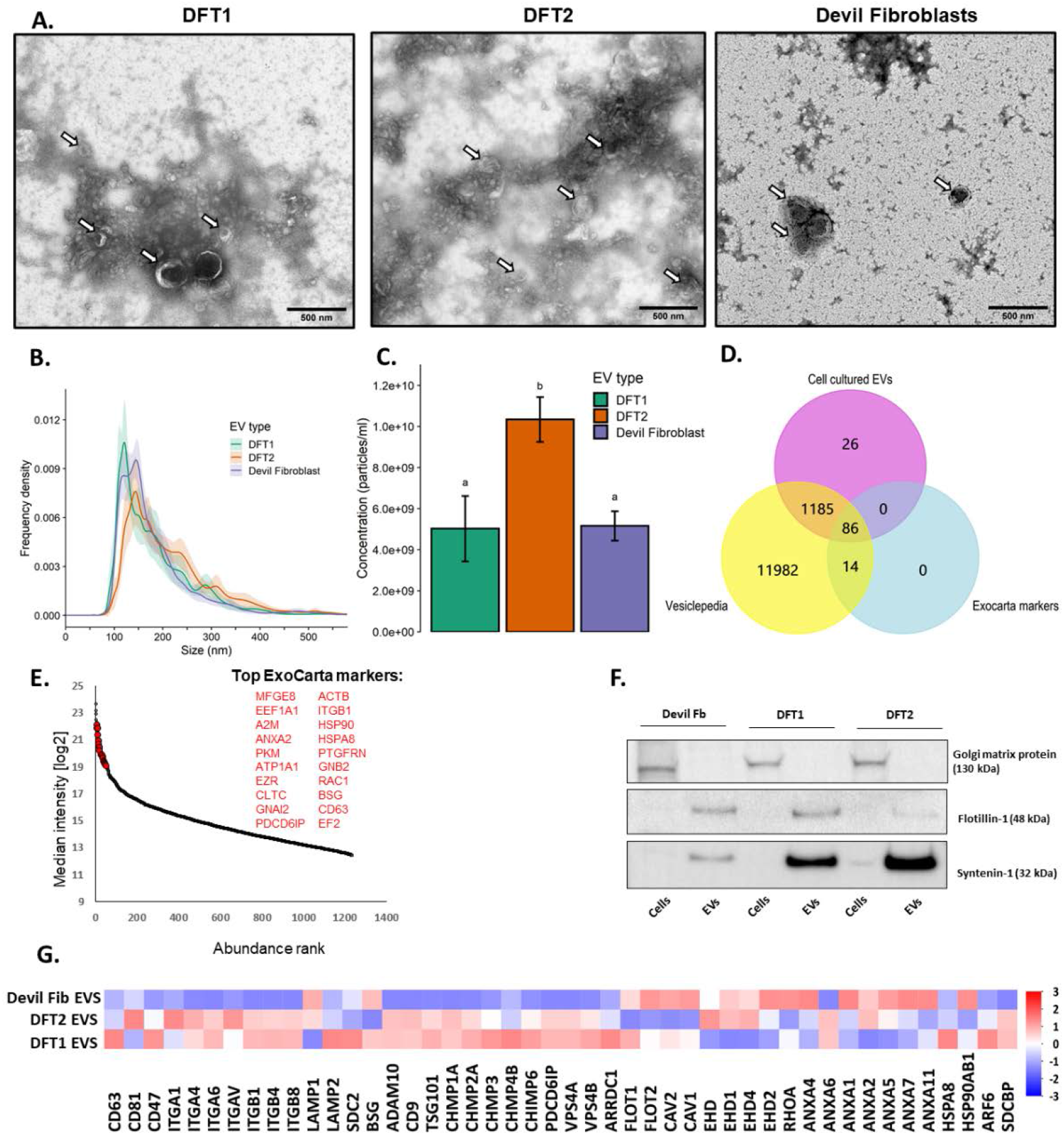
Characterization of cell cultured extracellular vesicles (EVs). **A.** Transmission electron microscopy images of isolated EVs from cell cultures. White arrows indicate EV structures. **B.** Particle size distributions of cell culture derived EVs measured by Nanoparticle tracking analysis (NTA), shaded areas represent 95% confidence intervals. **C.** NTA shows particle concentration and statistical significance is denoted by letters (ANOVA Tukey post-hoc test, p < 0.01), error bars represent 95% confidence intervals. **D.** Venn diagram of overlapping genes identified in EVs derived from cell cultures with Vesiclepedia, and the top one hundred exosomal genes reported in the Exocarta dataset. **E.** Median cell culture-derived EV protein abundance distribution as calculated from MS intensities of quantified peptides of each protein. The top twenty most abundant ExoCarta marker proteins are labelled in red. **F.** EV and cell lysates sample Western blots of a purity EV marker (Golgi matrix protein ~130 kDa) and two cytosolic EV markers (Flotillin-1 ~ 48 kDa and Syntenin-1 ~32 kDa). **G.** Heat map of patterns of mass spectrometry intensities of membrane and cytosolic EV markers in cell culture derived EVs. These EV markers are recommended by the MISEV 2018 guidelines (38). The bar color represents Z scores.

**Figure 1.**
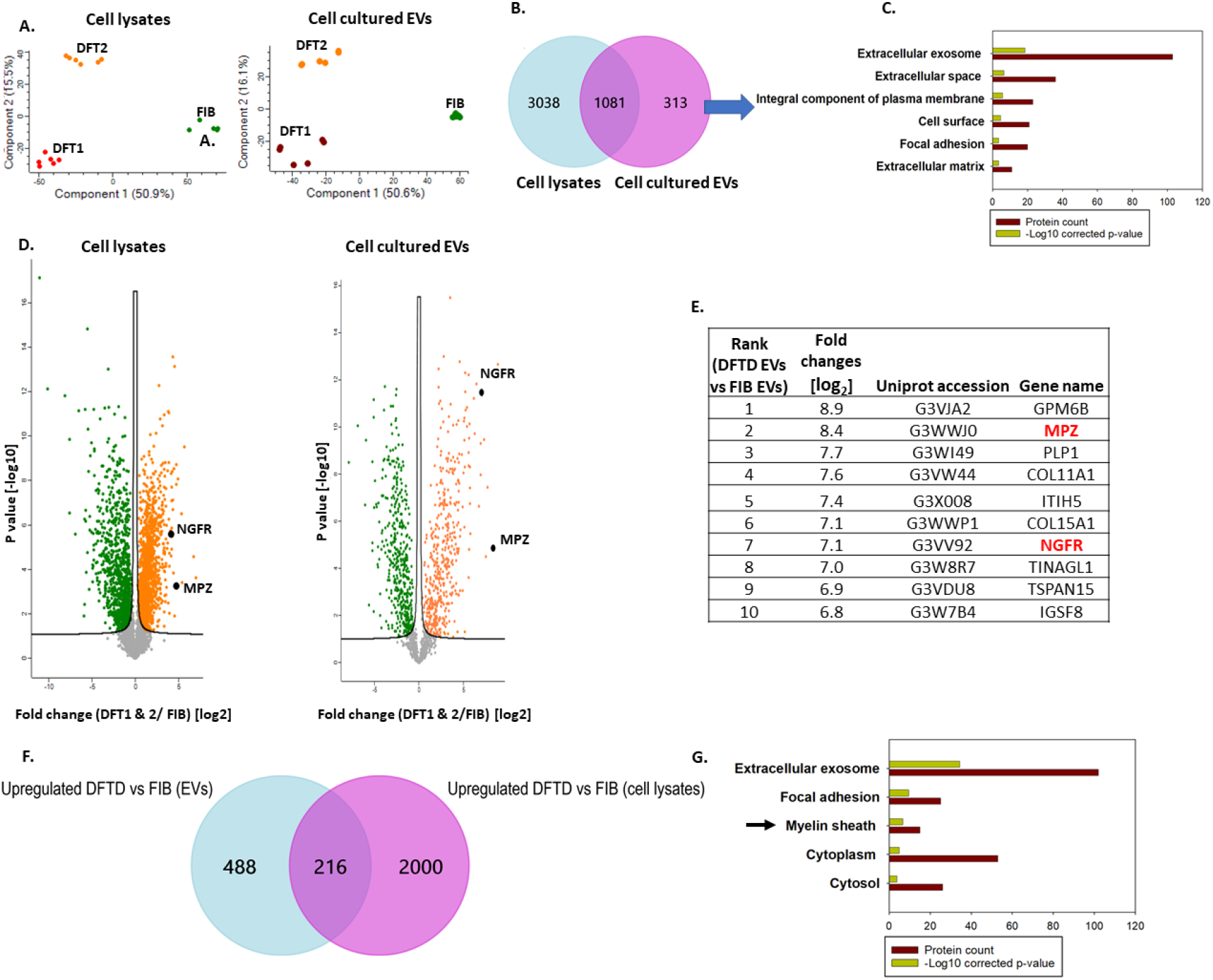
**A.** Principal component analysis (PCA) biplots of cell lysate and cell culture-derived EV proteomes. **B.** Venn diagram of cell lysate and cell culture-derived EV protein overlap. **C.** Over representing analysis (ORA) for cellular component gene ontology (GO) terms associated with proteins unique to cell cultured EVs (not present in cell lysates; 313 proteins). **D.** Volcano plots of log2 fold changes between DFTD (DFT1 and DFT2) and devil fibroblast cell lysates and cell culture derived EVs vs. fold change significance. Nerve growth factor and myelin protein zero are labelled in black. Orange denotes proteins significantly upregulated in DFTD cells EVs, green denotes proteins upregulated in devil fibroblast cells and EVs. Proteins above the black lines have a q-value < 0.05. **E.** Ten most-upregulated proteins in EVs derived from DFTD cells relative to EVs derived from devil fibroblasts. **F.** Venn diagram comparing upregulated proteins of DFTD EVs vs devil fibroblasts EVs and upregulated proteins of DFTD cells vs fibroblast cells. **G.** ORA analysis of cellular component GO terms associated with proteins that are upregulated in DFTD EVs relative to fibroblast EVs and also in DFTD cells relative to devil fibroblasts cells (216 proteins).

The post-filtering EV proteome from the DIA-MS analysis was converted to gene symbols for comparison with the human Vesiclepedia database. Gene symbols were retrieved for 1,297 proteins, while 97 proteins were uncharacterized, with no gene symbol reported. The EV proteome of devil samples shared 1,271 genes with the Vesiclepedia database (Fig. 1D). Of these Vesiclepedia-devil EV shared proteins, 86 proteins were also in the ExoCarta top-100 hundred proteins reported from EV preparations (Fig. 1D). The top-20 ExoCarta markers for EVs derived from cultured cells are plotted according to their abundance in the devil EV samples (Fig. 1E). These included several of the most common EV markers, PDCD6IP, CD63 and HSP90 (39). In addition to the proteins identified by mass spectrometry, western blotting was used for targeted analysis of the cytoplasmic EV markers Flotillin-1 (FLOT1) and Syntenin-1 (SDCBP) (Fig. 1F). These EV markers showed expression patterns that aligned with the corresponding proteomic data for these two proteins in Fig. 2E. The marker Golgi matrix protein (GM130) was absent in the Tasmanian devil EV samples, indicating purity in the EV preparation (38). Other membrane and cytosolic EV markers identified by mass spectrometry are represented as a heat map, which are recommended by the minimal information of extracellular 2018 guidelines (38) (Fig. 1G).

**Figure 2.**
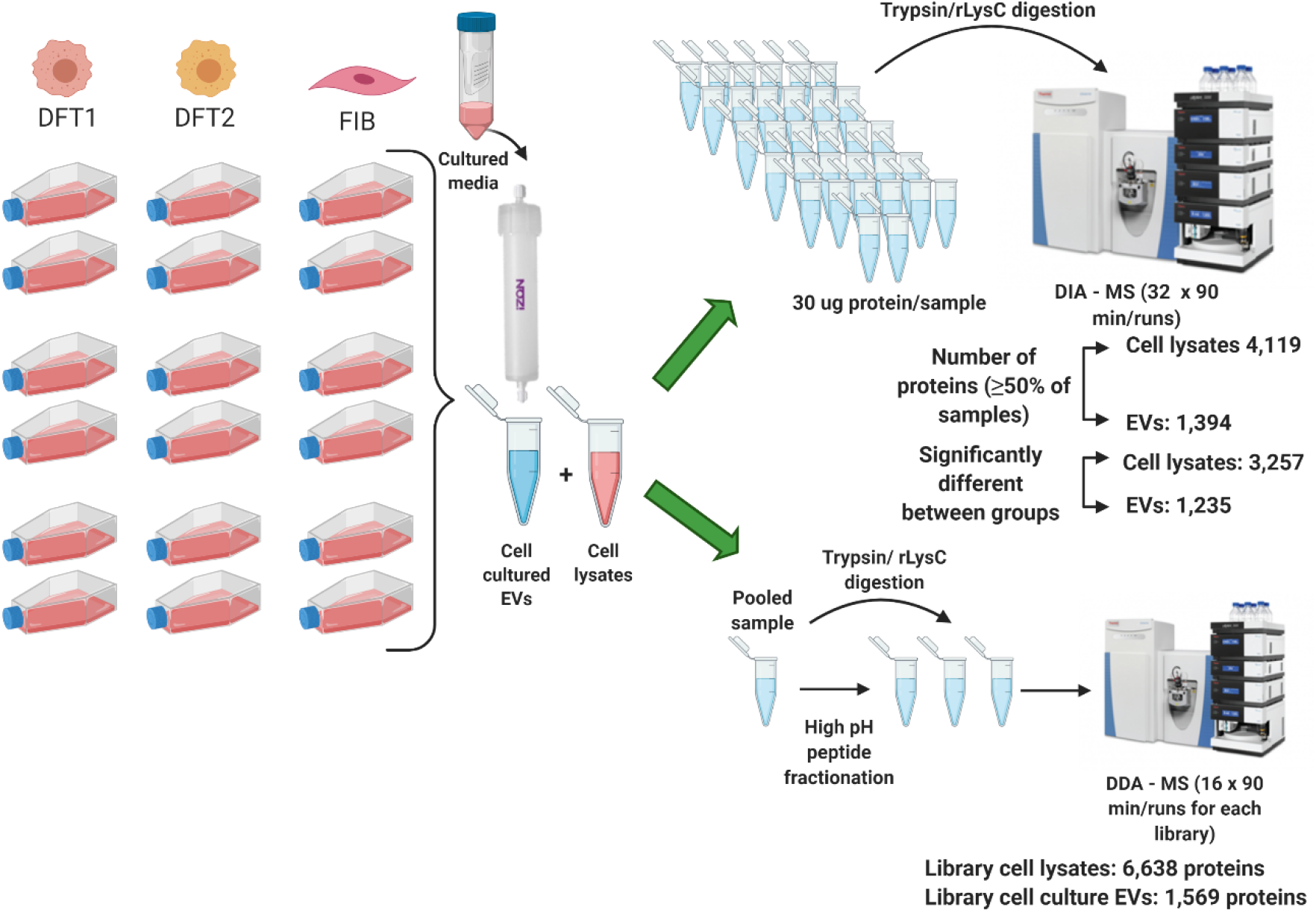
Schematic representation of the isolation of EVs derived from cell cultures. Proteomic workflow and an overview of mass spectrometry data is shown. The number of proteins from the two spectral libraries generated with dependent acquisition (DDA) mass spectrometry are reported in the figure. For the data independent acquisition (DIA) mass spectrometry, a total of 36 individual samples were analyzed, including 18 samples of cell lysates and 18 samples of cultured cell EVs. Significance of differences among the three types of cell lysate and cell culture-derived EV proteins are indicated at p < 0.05 and FDR < 0.05. Created with BioRender.com.

**Figure 2.**
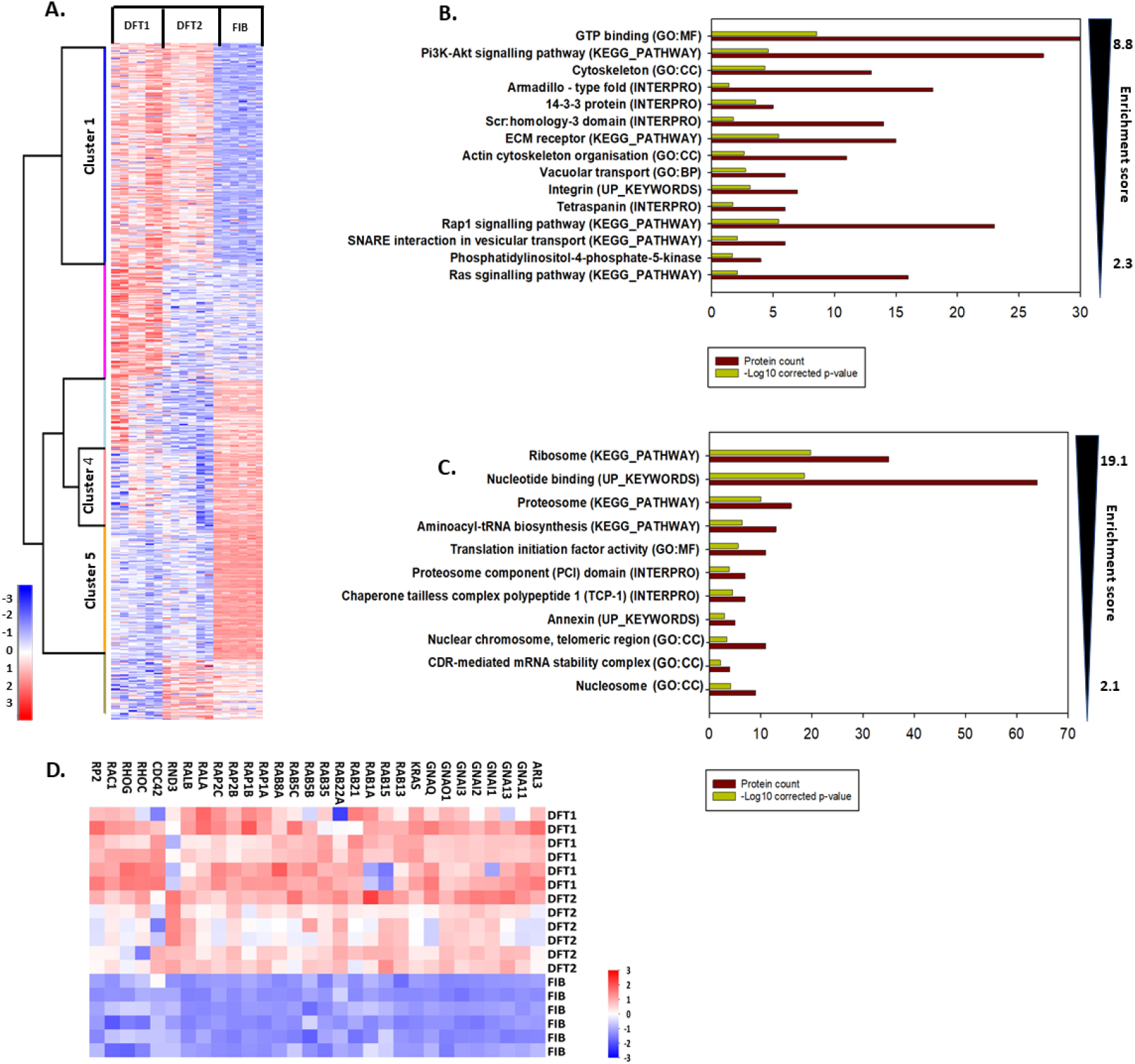
**A.** Hierarchical clustering of differential expressed DFTD (DFT1 and DFT2 combined) and devil fibroblast EV proteins with colour representing Z scores. **B.** Over representing analysis (ORA) for gene ontology (GO) terms, protein families and pathways of the upregulated proteins of DFTD (1 and 2) EVs (407 proteins), grouped in red color in cluster 1, and **C.** of the proteins upregulated in devil fibroblast EVs in clusters 4 and 5 (371 proteins). **D.** Expression of proteins belonging to the significantly enriched GTP binding GO term. The bar color represents Z scores.

### Proteomics experimental design and statistics overview

Spectral libraries were generated for cell lysates and EVs from cell cultures, based on DDA-MS of a pooled fractionated samples. The cell lysate library comprised 6,638 protein groups (from this point referred to as proteins), assembled from 80,845 stripped peptide sequences, while the EV library comprised 1,569 proteins, assembled from 15,705, stripped peptide sequences. Searching DIA-MS data for individual samples against their respective libraries identified 4,119 proteins in a minimum of 50% of the cell lysate samples and 1,394 proteins in a minimum of 50% of the EV samples. Comparison of DFT1, DFT2 and fibroblast protein intensity values by ANOVA (s0 = 0.1 and FDR = 0.05) identified 3,257 and 1,235 differentially-abundant proteins in cell lysate and EV samples, respectively (Table S6). A diagrammatic representation of the library generation and outcome is shown in Figure 2.

### EVs derived from DFTD cells represent their cell of origin

Principal component analysis (PCA) was first used to investigate patterns in proteomic composition of cell lysates and EVs from cultured cells within each sample type (Fig 3A). After filtering the DIA-MS data, variance of the 4,119 cell lysate proteins and 1,394 cell cultured EV proteins was explained by the first two principal components in each plot, respectively. For both the lysate and EV samples, PC 1 explained 50.9% and 50.6%, while PC 2 explained 15.5% and 16.1%, respectively. DFT1, DFT2 and fibroblast cell lysate and EV samples had a clear pattern, with replicates of each cell line type distinctly clustering together in the PCA plot.

**Figure 3.**
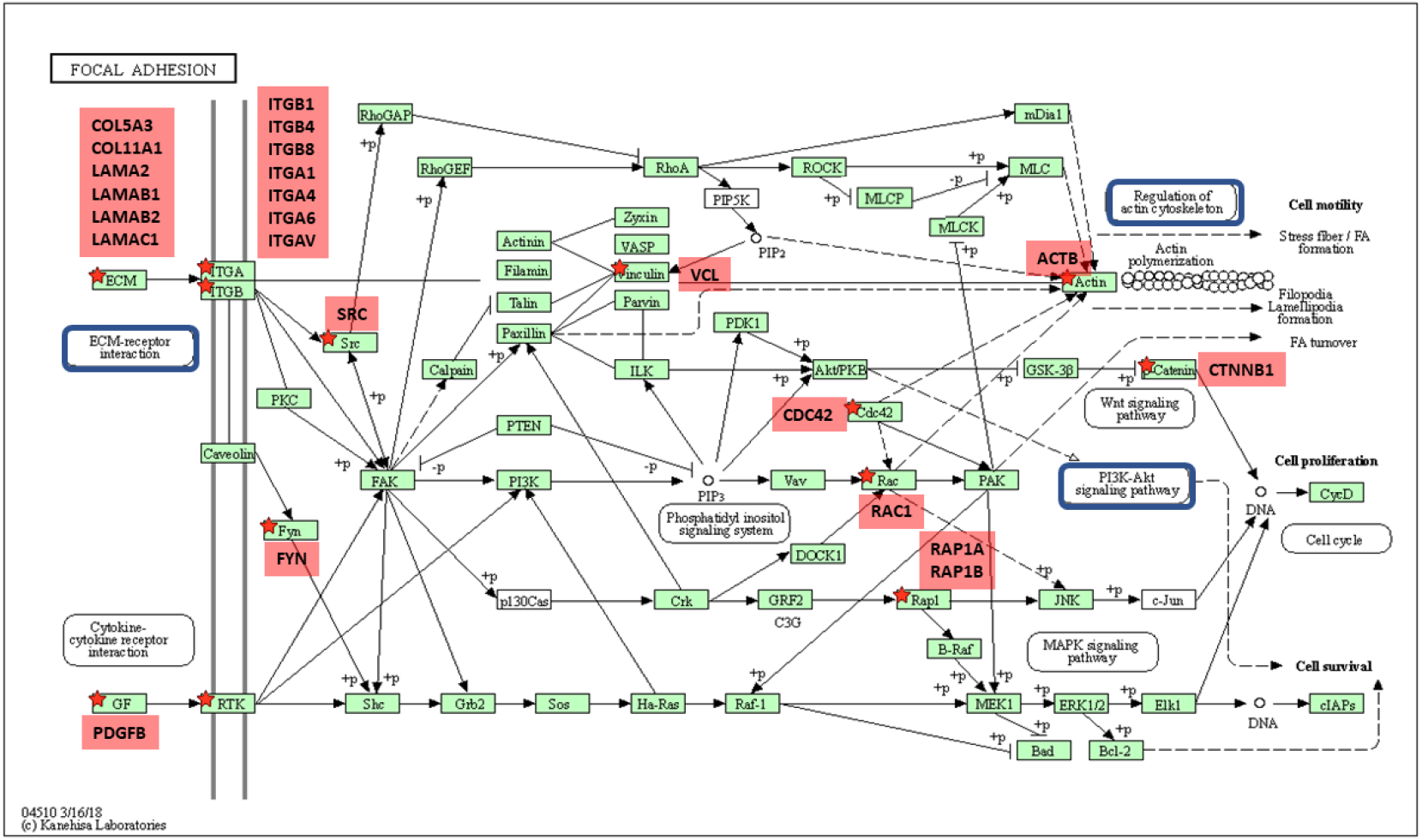
KEGG pathway map (shr04510) of Focal adhesion signaling pathway. Overview of interconnected signaling pathways and their genes names for the upregulated DFTD EV samples obtained from DAVID database. Additional KEGG pathways identified using DAVID analysis (ECM-receptor interaction, Regulation of actin cytoskeleton and PI3K-Akt signaling) are highlighted by the blue color. Proteins that are upregulated in DFTD EVs are highlighted by red stars and labelled with corresponding gene names.

To understand the relationship between sample types, we identified the proteins detected in EVs only, lysates only and in both datasets. Approximately three quarters of the EV proteins (1,081/1,394) were also detected in cell lysates (Fig. 3B). However, about one fifth of the EV proteins (313/1,394) were exclusively present in EVs (Fig 3B). Functional enrichment analysis of these only EV proteins revealed that the majority of them are annotated by the term extracellular exosome (Fig. 3C).

To determine whether EVs from DFTD cells reflect their cell of origin, the proteins upregulated in DFTD EVs and cells relative to devil fibroblasts EVs and cells were compared. Volcano plots show that DFTD EVs and cells expressed Nerve Growth Factor Receptor (NGFR) protein and Myelin Protein Zero (MPZ) in their top 1.2% percentile of upregulated proteins relative to devil fibroblast EVs and cells (Fig. 3D and 3E). Both proteins have been reported as highly expressed on DFT1 tumor biopsies, and have been used to diagnose DFTD tumors by immunohistochemistry (9). Moreover, of the EV proteins upregulated in DFTD relative to fibroblasts, 216 proteins were also upregulated in DFTD cell lysates relative to fibroblast lysates (Fig. 3F). These proteins showed enrichment of the gene ontology (GO) term Myelin Sheath (Fig. 3G), previously reported as a significant functional term associated with DFTD cells and biopsies (40).

### DFTD derived EVs enriched cell and focal adhesion proteins relative to fibroblasts derived EVs

To further characterize the protein signature of DFTD and devil fibroblast EVs, differentially abundant proteins between the three groups were identified using ANOVA (s0 = 0.1 and FDR = 0.05) and Z-scored proteomic data for the 1,235 significant proteins analyzed by hierarchical clustering. Proteomic data for DFT1, DFT2 and devil fibroblast derived EVs was displayed as a heatmap and the resulting tree was divided into 6 hierarchical clusters (Fig 4A). Cluster 1 contained 407 proteins upregulated in EVs derived from both cancers relative to EVs derived from devil fibroblasts (Table S7), while clusters 4 and 5 contained 371 proteins upregulated in EVs from devil fibroblasts compared to cancer EVs (Table S8).

**Figure 4.**
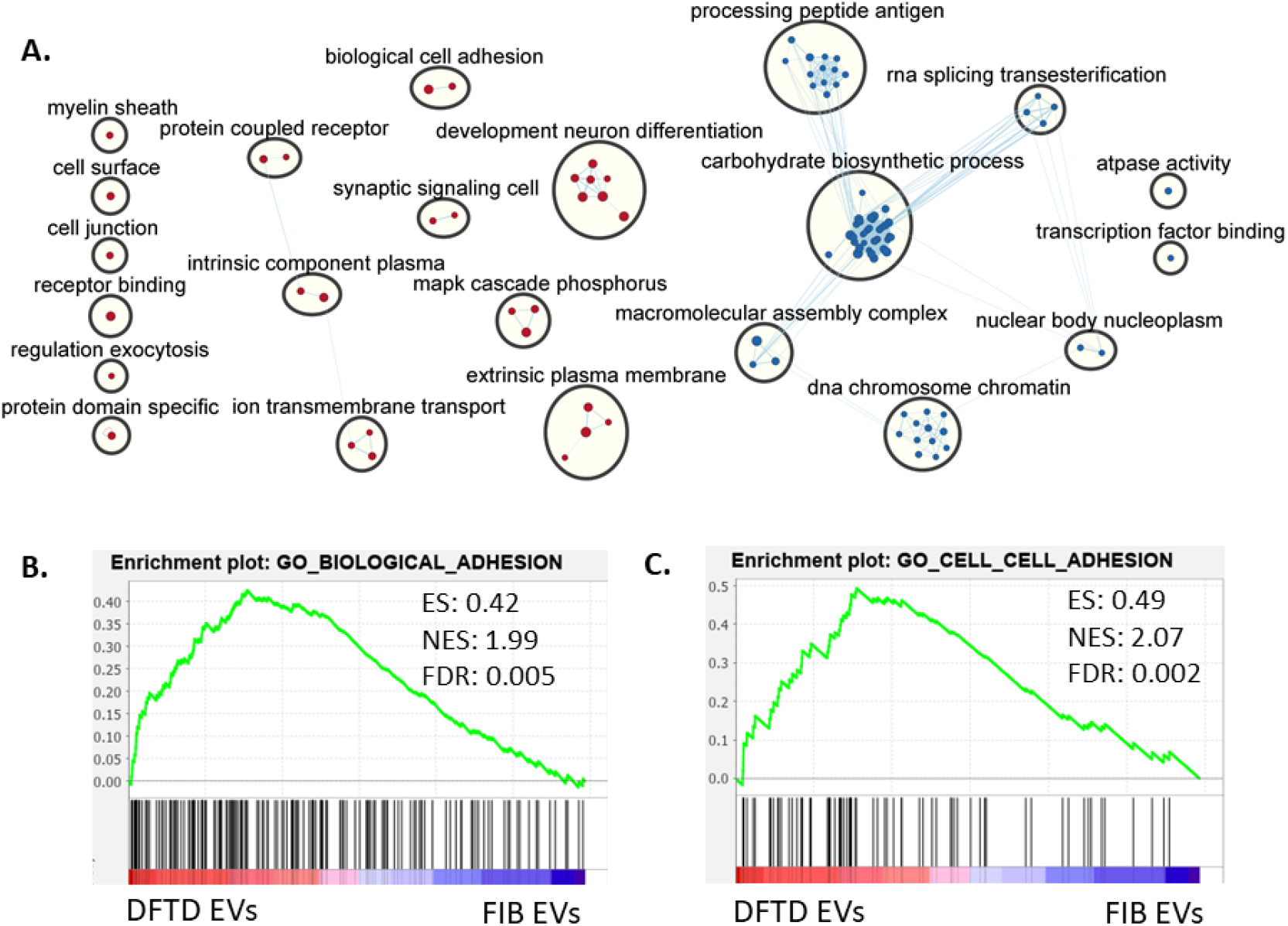
**A.** The enrichment map was created with parameter false discovery rate (FDR) q value <0.05, and combined coefficient >0.375 with combined constant = 0.5. Red and blue nodes represent EVs derived from DFTD cells and devil fibroblasts, respectively. The size of the oval increases as the gene set size is bigger. Gene set size varies from 185 to 25 genes. Normalized Enriched Score (NES) are from 2.3 to −3.23. **B.** Enrichment plots for the GO terms biological adhesion and **C.** cell-cell adhesion. Enrichments plot contains enrichment score (ES), normalized enrichment score (NES) and FDR q value. Both terms are included in the clustered term biological cell adhesion shown in Fig **6A**.

#### Over representation analysis (ORA)

The protein clusters identified above were then used for functional enrichment analysis, using over representation analysis (ORA) in DAVID software. ORA analysis of the DFTD-associated proteins revealed that many of the enriched functional terms were linked to focal adhesion (Fig 5), such as integrin, PI3K-Akt signalling pathway, ECM-receptor interaction, and regulation of actin cytoskeleton (Fig 4B). The highest enriched GO term for DFTD EVs was GTP binding, including RAS oncogenes and G-proteins alpha subunit which are involved in the RAP1 signaling pathway, also a significant KEGG pathway (41) (Fig 4.D). In contrast, proteins that were significantly more abundant in devil fibroblast EVs were enriched in functional terms related to protein biosynthesis such as ribosome, amino acyl tRNA biosynthesis and translation initiation, as well as protein folding (TCP-1), disposal (proteasome), and protein transport among others (Fig 4C).

**Figure 5.**
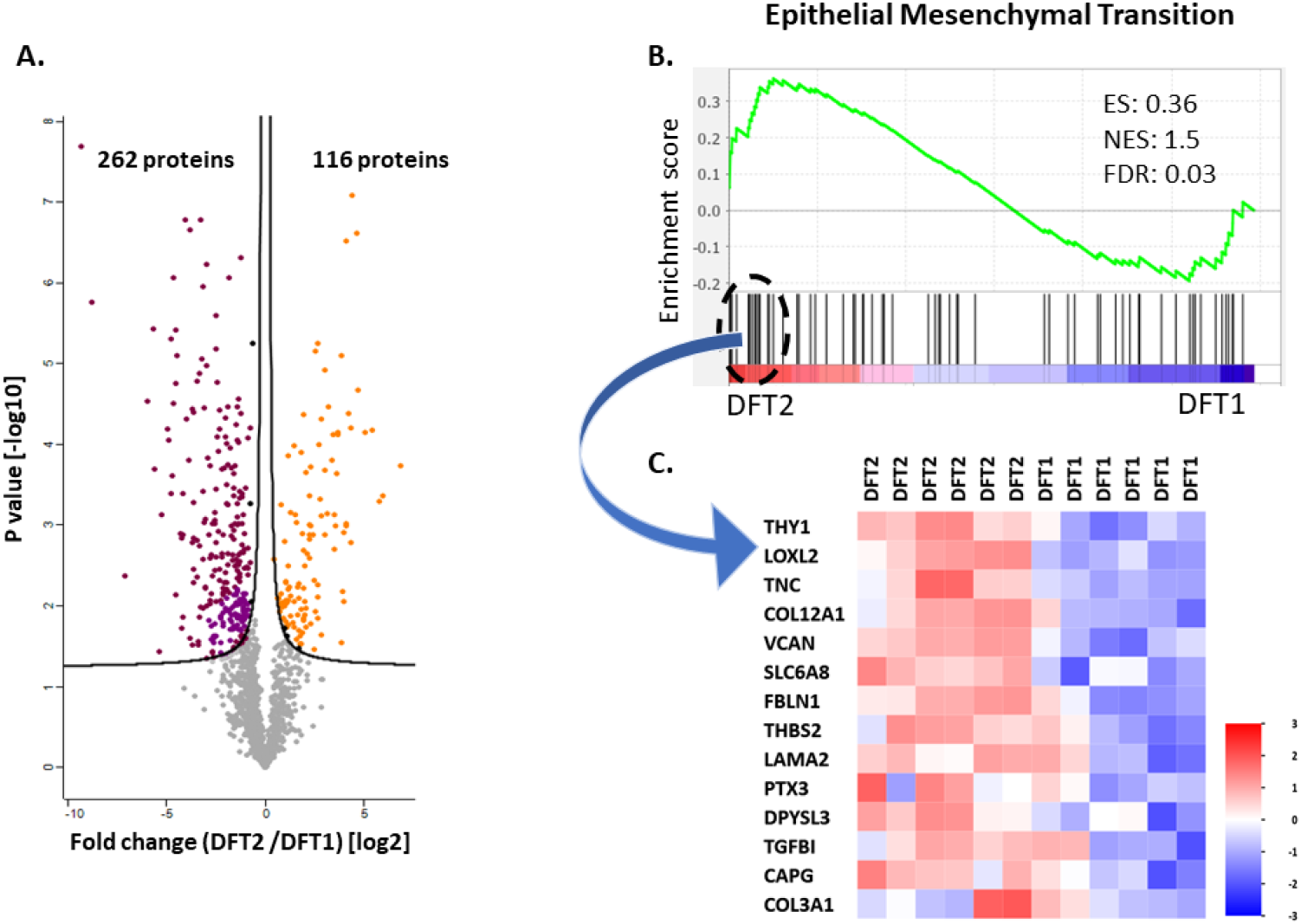
**A.** Volcano plot shows fold changes among proteome of EVs derived from DFT1 and DFT2 cells vs. statistical significance. Proteins labelled in orange are proteins upregulated in EVs derived from DFT2 cells (116), while proteins labelled in red are upregulated in EVs derived from DFT1 cells (262). **B.** Enrichment plot contains enrichment score (ES), normalized enrichment score (NES) and false discovery rate (FDR) q value. The bottom portion of the plot shows the genes belonging to the hallmark, and they are ranked according to their differential expression. Higher and lower expression are represented by red and blue color, respectively. **C.** Heat map showing the core list of proteins that contributes the most to the Epithelial Mesenchymal Transition. The bar color represents Z scores.

#### Gene enrichment set analysis (GSEA)

To further distil biological information from our proteomics data, the complementary GSEA approach identified enrichment of protein groups in DFTD derived EVs relative to fibroblast derived EVs associated with biological cell adhesion, MAP kinase cascade, development neuron differentiation, synaptic signaling and others (Fig 6A). In contrast, GSEA revealed enrichment of protein groups in devil fibroblasts derived EVs including carbohydrate biosynthetic process, processing peptide antigen, macromolecular assembly complex, and others (Fig 6A).

### DFT2 derived EVs enriched the epithelial mesenchymal transition hallmark relative to DFT1 derived EVs

In addition to using the DFT1, DFT2 and fibroblast data to identify a tumor EV-associated signature, we could also identify differentially abundant proteins between EVs derived from DFT2 and DFT1 cells. Based on t-test analyses, 116 and 262 proteins, (q-value < 0.05) were up-regulated in DFT2 and DFT1 EVs, respectively (Fig.7A). Comparison of the EV datasets from DFT1 and DFT2 by GSEA showed that EVs derived from DFT2 cells were significantly enriched in the hallmark epithelial mesenchymal transition proteins relative to EVs derived from DFT1 cells (normalized enrichment score = 1.5 and FDR = 0.03) (Fig 7B).

## Discussion

Despite metastasis being a primary cause of Tasmanian devil death due to DFTD (8), attempts to uncover the mechanistic basis of metastasis in DFTD have been limited. This is largely due to challenges inherent to collecting samples from wild animals, such as the logistical difficulty of capturing animals and the detrimental effects of invasive biopsies. Here, we have demonstrated the potential utility of EVs to better understand DFTD metastasis. Furthermore, these results provide a basis for future analysis of EVs derived from Tasmanian devil biofluids to investigate DFTD pathogenesis *in vivo*.

Consistent with the Schwann cell origin of DFTD (2, 6, 40), DFTD EVs displayed the myelin proteins NGFR and MPZ which are used as diagnostic markers for DFTD on tumor biopsies (9). Further signatures of Schwann cell origin including the myelin sheath GO term and clusters of neuronal and glial function-related terms were enriched in the DFTD EV proteome. The maintenance of key cell of origin signatures has been reported in EVs obtained from other cancer cell lines (42–44). Together, these findings provide evidence that DFTD EVs retain hallmarks of their cancer cell origin and suggest that tumor derived EVs could be identified in devil body fluid samples.

The functional pathways enriched in the DFTD EV proteome suggest that EVs may be involved in metastatic processes in the progression of both cancers. The focal adhesion pathway is formed by large protein complexes which are upregulated in cancer cells to colonize other organs during metastasis (45). Several of the cell adhesion-related proteins highly enriched in DFTD EVs have been implicated in EV-associated metastasis, specifically in preparing the pre-metastatic niche in the lungs. In particular, the ras-related proteins Ra1A and Ra1B, shown to promote lung metastasis in breast cancer (46) were upregulated in both DFT1 and DFT2 EVs relative to fibroblast EVs. We also found that the integrin subunit ITGA6, which can moderate tumor EV organotropism to the lung (20), was also significantly upregulated. Considering the role of these proteins in lung-specific metastasis together with the finding that nearly 50% of tumor metastasis in DFTD involves the lungs (5), our findings raise the possibility that DFTD EVs play a role in preparing the pre-metastatic niche in the lung tissue of infected devils.

Differences in the EV proteomes of DFT1 and DFT2 cells suggest a mechanistic basis of a potential increased of aggressiveness in DFT2. The epithelial mesenchymal transition (EMT) processes, differentially enriched in DFT2 EVs, are known to confer mesenchymal properties that increase cell motility and migration (47). As such the EMT hallmark is linked to increased aggression cancer phenotypes and metastatic behavior (48-51). The EMT characteristics of the DFT2 EV proteome are consistent with a ‘repair’ Schwann cell phenotype, the transition into which involves the re-activation of EMT related processes (52). DFT2 tumor biopsies demonstrated the repair Schwann cell phenotype relative to peripheral nerves, while DFT1 biopsies did not show significant enrichment of this phenotype (40). Given this combined evidence, we propose that the EMT traits of DFT2 EVs could reveal clues about DFT2 tumorigenesis and possibly indicate more significant metastatic behavior. Further, the differential expression of EMT hallmark proteins offers potential for differential diagnosis between DFT1 and DFT2.

The development of standardized methods for isolation of EVs of sufficient yield and purity, especially from body fluids, is an ongoing research activity. EVs from body fluids are derived from diverse cell populations and usually are co-isolated with highly abundant contaminants present in body fluids such as albumins and lipoproteins, which can lead to erroneous interpretation in downstream analyses (53). Thus, the use of cell cultures enables the study of EVs derived from a single and well-defined cell type and the ability to manipulate conditions to reduce these body fluid contaminants (54). Further, we were able to optimize coverage of EV proteins in this study with DIA-MS techniques and targeted data analysis using EV protein spectral libraries (55, 56). However, our findings must be considered in light of the challenges to extrapolate results from *in vitro* to *in vivo* models, and also corroborating them using functional assays. Although cultured cell lines are extremely valuable for research on cancer biology and metastatic mechanism, they cannot completely recapitulate the complexity of cancer in an organism (57). For this reason, a necessary next step is an investigation of the EV composition of biofluids of DFTD-affected devils.

This is the first investigation, to our knowledge, of EVs in the context of a cancer that affects wild animals. We have demonstrated the potential for EVs to shed light on mechanisms of DFTD metastasis, as well as identify novel candidate proteins with potential value as diagnostic and prognostic biomarkers in devil biofluid samples. Metastatic cancers have been increasingly reported in wildlife in the past few decades (58). EV approaches offer a promising avenue to the development of sensitive and less-invasive clinical tools needed for wildlife cancer monitoring and management. The EV-based investigation of cancer in the wild will likely provide useful information for human cancers, as EVs are well-conserved structures through the tree of life (59), and wild animals have more similarities to humans than laboratory animals in terms of environmental exposures and life-span (60).

## Acknowledgements

The authors would like to acknowledge all the members of the devil and wild immunology group for their advice and guidance. We would also like to thank the School of Medicine at the University of Tasmania for providing support for this research. This work was supported by the National Geographic explorer early career grant, Holsworth Wildlife Research Endowment grant, the University of Tasmania Foundation through funds raised by the Save the Tasmanian Devil Appeal, and the Tasmanian School of Medicine 2020 HDR student funding. Proteomics infrastructure was funded by ARC LE180100059.

## Data availability

The mass spectrometry raw proteomics data have been deposited to the ProteomeXchange Consortium via the PRIDE (61) partner repository with the dataset identifier PXD020766 (62).

