## Supplemental material for "Tasmanian devil facial tumor-derived extracellular vesicles reveal mesenchymal transition markers and adhesion molecules related to metastasis"

**Title**

**Supplementary data**

Table S1. Description of cell lines used in this study.

Table S2. Description of antibodies used in this study.

**Supplementary excel file 1. (EV summary):**

Table S3. Spectronaut quant pivot report with number of peptides included.

Table S4. EV proteins before filtering and non-imputed.

Table S5. EV proteins after filtering and non-imputed.

Table S6. EV proteins after filtering and imputed.

Table S7. EV proteins (imputed) that are significantly different between groups. Fold changes between DFTD EV proteins vs Fibroblast EV proteins. Fold changes between DFT1 EV proteins vs. DFT2 EV proteins.

Table S8. DFTD EV proteins upregulated vs. fibroblast EV proteins (imputed) included in cluster 1 from the hierarchical cluster analysis.

Table S9. Fibroblast EV proteins upregulated vs. DFTD EV proteins (imputed) included in cluster 4 and 5 from the hierarchical cluster analysis.

**Supplementary excel file 2. (Cell lysate summary):**

Table S11. Spectronaut quant pivot report with number of peptides included.

Table S10. Cell lysate proteins before filtering and non-imputed.

Table S11. Cell lysate proteins after filtering and non-imputed.

Table S12. Cell lysate proteins after filtering and imputed.

Table S13. Cell lysate proteins (imputed) that are significantly different between groups. Fold changes between DFTD cell lysate proteins vs Fibroblast cell lysates proteins. Fold changes between DFT1 cell lysate proteins vs DFT2 cell lysate proteins.

**Supplementary text file 3. Setup file for DIA analysis of EVs.**

**Supplementary text file 4. Setup file for DIA analysis of cell lysate.**

| | **Tasmanian devil microchip number** | **Location** | **DFTD strain** |
|---|---|---|---|
| DFT1 cell line: C5065 | 982009100322308 | Bangor | Strain 3 |
| DFT1 cell line: 4906 | 982009100873958 | Coles Bay | Strain 4 |
| DFT1 cell line: 1426 | 985120016021248 | Fentonbury | Strain 2 |
| DFT2 cell lines: RV | 982000190608331 | Cygnet | N/A |
| DFT2 cell lines: Snug | 982000356526917 | Lower Snug | N/A |
| DFT2 cell lines: Jarvis | 982000356669303 | Snug Tiers | N/A |
| Fibroblasts: Lucy | 982000167813072.00 | Richmond | N/A |
| Fibroblasts: Rosie | 982000167871303.00 | Richmond | N/A |
| Fibroblasts: Stanley | 982009105160330.00 | Richmond | N/A |

Table S1. Description of cell lines used in this study. Location refers to the place in which the devil was trapped and sampled.

| **Antibody** | **Manufacturer** | **Catalog #** | **Clone** | **Species** | **Immunogen** |
|---|---|---|---|---|---|
| Anti-Flotillin-1 | Biosciences | 610821 | Clone 18 | Mouse IgG1 | Mouse Flotillin aa. 312-428 |
| Anti-Syntenin | abcam | ab19903 | Polyclonal | Rabbit Polyclonal | Synthetic peptide corresponding to Mouse Syntenin aa 250 to the C-terminus (C terminal) conjugated to keyhole limpet hemocyanin. |
| Anti-GM130 | Biosciences | 610822 | Clone 35 | Mouse IgG1, K | Rat GM130 aa. 869-982 |

Table S2. Description of cross-reactive antibodies used in this study.
